# Direct ETTIN-auxin interaction controls chromatin state in gynoecium development

**DOI:** 10.1101/863134

**Authors:** André Kuhn, Sigurd Ramans Harborough, Heather M. McLaughlin, Stefan Kepinski, Lars Østergaard

**Author notes:** For correspondence, Mailing address of corresponding author Lars Østergaard, John Innes Centre Norwich Research Park Norwich, NR4 7UH, United Kingdom, Skype: lars.oest. Impact statement: Auxin binds to the ETTIN transcription factor to disrupt the interaction between ETT and a TPL/TPR co-repressor and subsequently affecting chromatin dynamics to ensure proper gynoecium development.

## Abstract

Hormonal signalling in animals often involves direct transcription factor-hormone interactions that modulate gene expression^1, 2^. In contrast, plant hormone signalling is most commonly based on de-repression via the degradation of transcriptional repressors^3^. Recently, we uncovered a non-canonical signalling mechanism for the plant hormone auxin in organ development with strong similarity to animal hormonal pathways. In this mechanism, auxin directly affects the activity of the auxin response factor ETTIN (ETT) towards regulation of target genes without the requirement for protein degradation^4, 5^. Here we show that auxin binds ETT to modulate gene expression and that this ETT-auxin interaction leads to the dissociation of ETT from co-repressor proteins of the TOPLESS/TOPLESS-RELATED family followed by histone acetylation and the induction of target gene expression. Whilst canonical ARFs are classified as activators or repressors^6^, ETT is able to switch chromatin locally between repressive and de-repressive states in an instantly-reversible auxin-dependent manner.

---

Developmental programmes within multicellular organisms originate from a single cell (*i.e.* a fertilised oocyte) that proliferates into numerous cells ultimately differentiating to make up specialised tissues and organs. Tight temporal and spatial regulation of the genes involved in these processes is essential for proper development of the organism. Changes in gene expression are often controlled by mobile signals that translate positional information into cell-type specific transcriptional outputs^7^. In plants, this coordination can be facilitated by phytohormones such as auxin, which controls processes throughout plant development^8^. In canonical auxin signalling, auxin-responsive genes are repressed when auxin levels are low by Aux/IAA transcriptional repressors that interact with DNA-bound Auxin Response Factors (ARFs). As auxin levels increase, the auxin molecule binds to members of the TIR1/AFB family of auxin co-receptors^9, 10^. This facilitates interaction with Aux/IAA repressors, Aux/IAA ubiquitinylation and subsequent degradation by the 26S proteasome, while relieving the repression of ARF-targeted loci^11, 12^.

We recently identified an alternative auxin-signalling mechanism whereby auxin directly affects the activity of a transcription factor (TF) complex towards its downstream targets^4, 5^. This mechanism mediates precise polarity switches during organ initiation and patterning and includes the ARF, ETTIN (ETT/ARF3) as a pivotal component. However, ETT is an unusual ARF lacking the Aux/IAA-interacting Phox/Bem1 (PB1) domain^4, 13^ and it is therefore likely that ETT would mediate auxin signalling via an alternative pathway.

ETT can interact with a diverse set of TFs and these interactions are sensitive to the naturally occurring auxin, indole 3-acetic acid (IAA). The region responsible for IAA-sensitivity is situated within the C-terminal part of ETT, known as the ETT-Specific (ES) domain^6^. A protein fragment containing 207 amino acids of the ES domain, ES^388–594^, sufficient for mediating IAA-sensitivity in ETT-protein interactions, was produced recombinantly and shown to be intrinsically disordered^14^. The sensitivity of ETT-TF interactions to IAA suggests a direct effect of the IAA molecule on the ETT protein.

Therefore, to test whether ETT binds IAA, we carried out heteronuclear single quantum coherence (HSQC) nuclear magnetic resonance (NMR) experiments using ^15^N-labelled ES^388–594^ protein. The HSQC spectrum, recorded at 5°C, shows a prominent signal-dense region consistent with the ES domain being largely intrinsically disordered. Interestingly, the spectrum also shows dispersed peaks flanking the signal-dense region indicating that there is nevertheless some propensity to form secondary structure, particularly with a helical character (Fig. 1a). In addition to this overview of ETT structure, the HSQC NMR probes chemical shifts of protein amide-NH bonds in response to the presence of ligand^15^. We found that a number of residues shifted their position in the spectrum in response to the addition of IAA, whereas addition of the related Benzoic Acid (BA) had no effect (Fig. 1a-c). These shifts show that certain residues are experiencing a changed chemical environment as a consequence of IAA-binding and this may include the conformational change of a structural motif within the ETT protein. The HSQC experiment therefore demonstrates that ETT binds IAA directly. This experiment has not allowed us to assign signals to specific amino acids and hence there is some uncertainty associated with tracking the chemical shifts of some residues. However, a particularly large change is observed when IAA is added to the ETT fragment for the tryptophan NH cross peak (∼10ppm, rectangle I in Fig. 1a,c). Since there is only one tryptophan in the ETT fragment used here (W505), this shift can be assigned to this residue.

**Figure 1.**
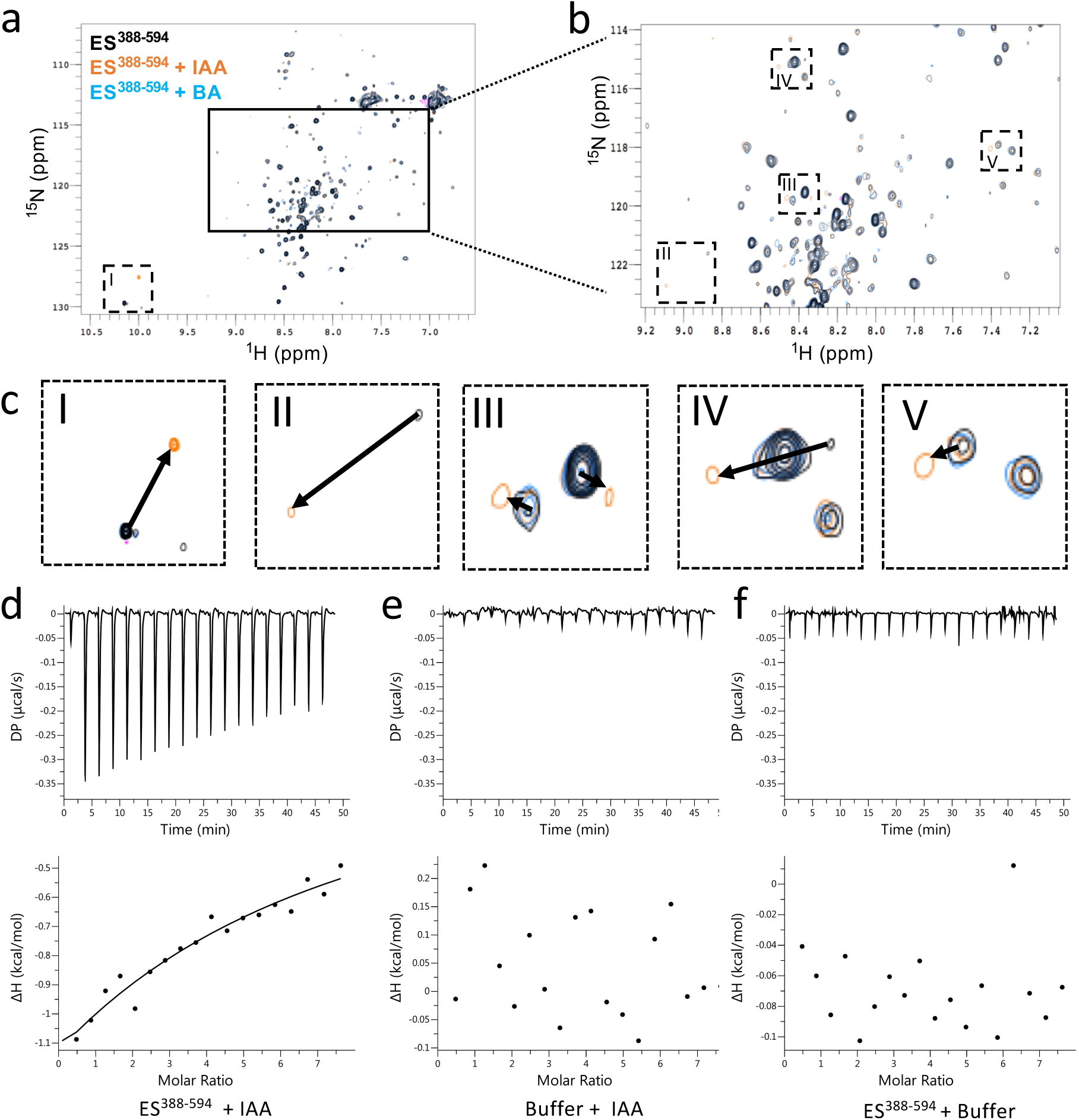
ETT directly binds auxin (IAA). **a,** HSQC-NMR performed with ES^388-594^ protein either alone (black), with indole-3-acetic acid (IAA, orange) or benzoic acid (BA, blue). **b,** zoom-in of the indicated rectangular region in a. **c,** zoom-in of the specific shifts (labelled I-V) in the indicated dotted rectangles in a and b. Changes in chemical shifts are indicated by arrows from control to IAA treatment. **d-f,** ITC spectre showing heat exchange between ES^388-594^ protein and IAA (**d**), but not in controls (**e,f**). See Figure 1-figure supplement 1 for parameters used in the HSQC-NMR experiment.

We also used the recombinant ETT fragment in an Isothermal Titration Calorimetry (ITC) assay, which characterises binding of ligands to proteins by determining thermodynamic parameters of the interaction as heat exchange. This experiment revealed interaction between ETT and IAA, while control experiments titrating IAA into buffer without protein and titrating buffer without IAA into the ETT fragment showed no heat exchange (Fig. 1d-f). Together, these two independent biochemical methods demonstrate that ETT binds IAA directly thus revealing a key molecular aspect of the non-canonical auxin-signalling pathway.

Previously, *PINOID* (*PID*)^16^ and *HECATE1* (*HEC1*)^17^ were identified as ETT target genes^4, 5^ and both genes are upregulated in gynoecium tissue from the *ett-3* mutant compared to wild type (Figure 2-figure supplement 1). We also observed that expression of both genes is induced by IAA, but did not observe any additional induction beyond the constitutive upregulation in the *ett-3* mutant background (Figure 2-figure supplement 1). This ETT-dependent regulation does not require a functional TIR1/AFB machinery, since IAA-induction of *PID* and *HEC1* is still observed in TIR1/AFB mutant combinations, whereas the known TIR1/AFB-mediated auxin induction of the *IAA19* gene is completely abolished in these mutants (Fig. 2a-c).

**Figure 2.**
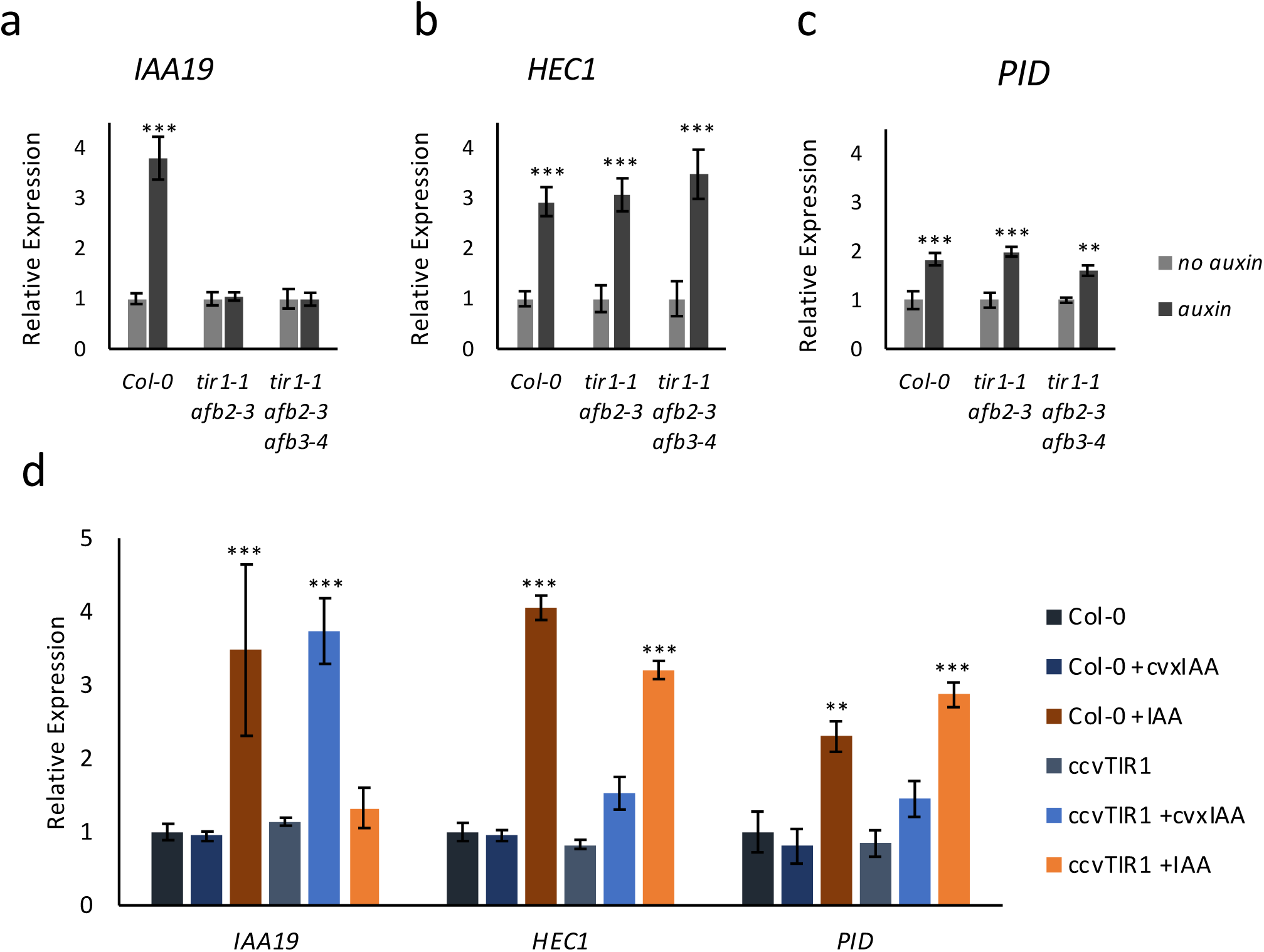
ETT regulates target gene expression independently of TIR1/AFB auxin receptors. Expression of the canonical auxin responsive *IAA19* gene (**a**) and the ETT-target genes *HEC1* (**b**) and *PID* (**c**) in control-treated or 100 µM IAA-treated gynoecia assayed using qRT-PCR. **a,** *IAA19* expression is up-regulated in response to auxin in wild-type gynoecia (*Col-0*) but not in *tir1/afb* double and triple mutants. The ETT-target genes *HEC1* and *PID* are up-regulated in response to auxin in both wild-type and auxin receptor mutants (**b, c**). This suggests a TIR1/AFB independent regulation of these genes. **d,** Expression of *IAA19*, *HEC1* and *PID* in response to treatment with 100 µM IAA and 100 µM cvxIAA in wild-type (*Col-0*) and *pTIR1:ccvTIR1* gynoecia in the *tir1 afb2* double mutant (*ccvTIR1*). The data confirm TIR1/AFB independent regulation of *HEC1* and *PID* in the gynoecium. ***p <0.0001; Shown are mean ± standard deviation of three biological replicates. See Figure 2-source data 1 for statistical analyses.

To further assess the TIR1/AFB independence of the ETT-mediated auxin signalling pathway, we exploited a recently-developed synthetic auxin-TIR1 pair^18^. In this system, the auxin-binding pocket of TIR1 has been engineered (ccvTIR1) to accommodate an IAA derivative bearing a bulky side chain (cvxIAA). By expressing the ccvTIR1 in a *tir1 afb2* mutant background, the canonical pathway will only respond to the addition of cvxIAA and not IAA^18^. We performed an expression analysis on *ccvTIR1* gynoecia treated ±cvxIAA and ±IAA as well as control plants with the same treatments. In this experiment, *IAA19* served as a control gene whose expression is known to be regulated in a TIR1/AFB-dependent manner. Indeed, *IAA19* was strongly upregulated by cvxIAA in the ccvTIR1 line, but not by IAA (Fig. 2d). In contrast, *PID* and *HEC1* expression was not significantly affected by cvxIAA, whilst still responding to IAA in the ccvTIR1 background (Fig. 2d). These data demonstrate that ETT-mediated auxin signalling can occur independently of the canonical TIR1/AFB signalling pathway.

In a phylogenetic analysis of ETT protein sequences across the angiosperm phylum, we identified a number of regions that are highly conserved (Figure 3-figure supplement 1). Unsurprisingly, the DNA-binding domain characteristic to B3-type TFs such as ARF proteins was conserved across all ETT proteins. Towards the C terminus of the ES domain we identified an EAR-like motif with a particularly high level of conservation (Fig. 3a, Figure 3-figure supplement 1). Ethylene-responsive element binding factor-associated Amphiphilic Repression (EAR) motifs are also found in Aux/IAA proteins. Interactions between Aux/IAA and members of the TOPLESS and TOPLESS-RELATED (TPL/TPR) family of co-repressors occur via this motif^19^. TPL/TPRs mediate their repressive effect by attracting histone deacetylases (HDACs) to promote chromatin condensation^20^. Since ETT functions independently of the canonical auxin pathway, it is possible that its role in chromatin remodelling occurs via direct interaction with TPL/TPRs through the EAR-like motif. To test this, we carried out Yeast 2-Hybrid (Y2H) assays in which ETT was found to interact with TPL, TPR2 and TPR4 (Fig. 3b, Figure 3-figure supplement 1). Moreover, mutating residues in the EAR-like motif abolished the interactions demonstrating its requirement for the ETT-TPL/TPR interaction (Fig. 3b).

**Figure 3.**
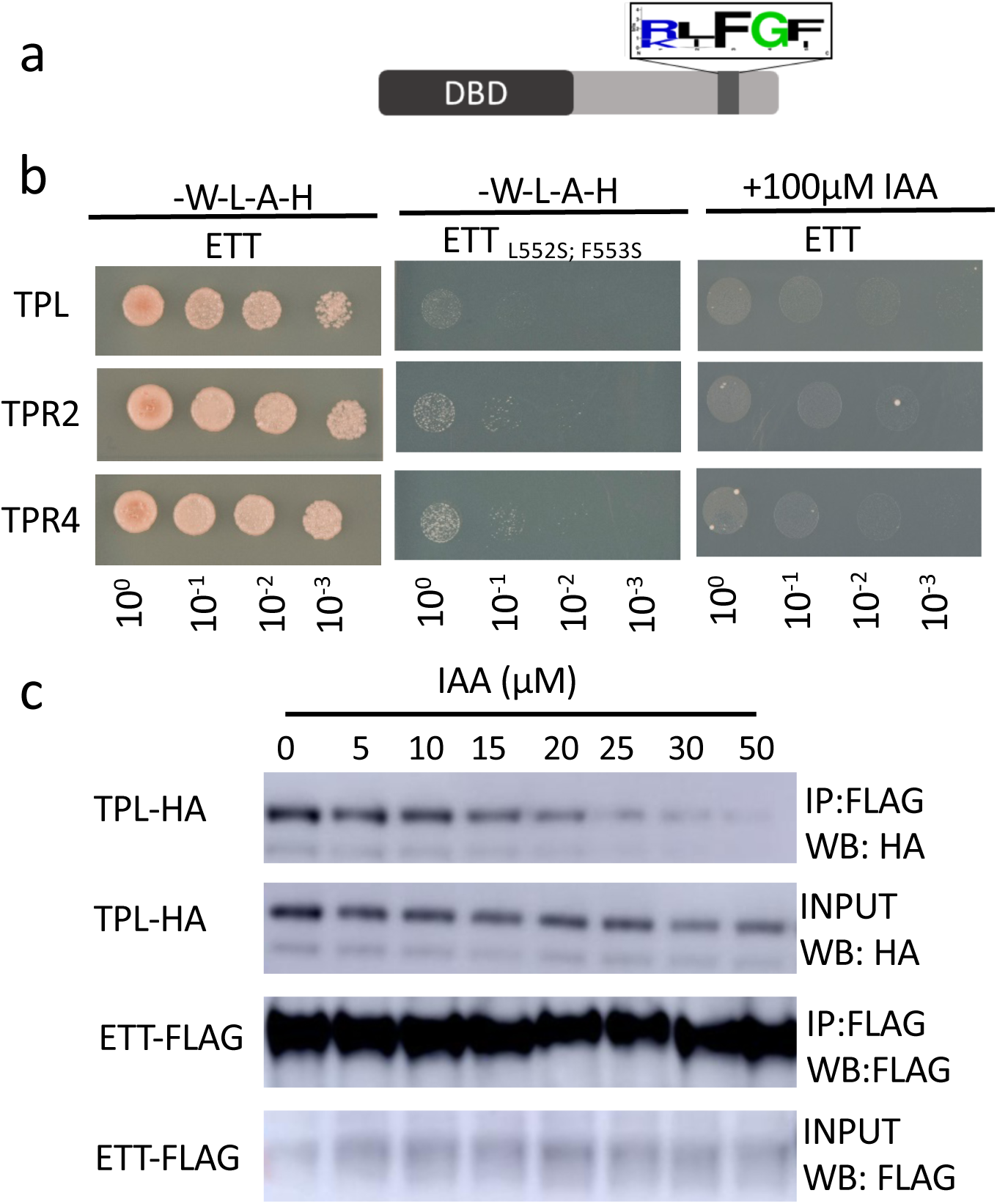
ETT interacts with members of the TPL/TPR co-repressor family in an auxin-sensitive manner. **a,** Schematic representation of ETT protein highlighting an EAR-like motif in the C-terminal ETT-specific domain. **b,** Y2H showing that ETT interacts with TPL, TPR2 and TPR4. These interactions depend on the identified C-terminal RLFGF motif and are auxin-sensitive. DBD, DNA-binding domain. **c,** Co-IP revealing that ETT interacts with TPL in an auxin-sensitive manner with increasing IAA concentrations weakening the interaction.

Given that several ETT-protein interactions are affected by IAA and that part of the ETT transcriptome changes in response to IAA^4, 5^, we tested the IAA sensitivity of ETT-TPL/TPR interactions. In both Y2H and in co-immunoprecipitation (Co-IP) experiments, we observed that the interactions were reduced with increasing IAA concentrations (Fig. 3b,c and Figure 3-figure supplement 2). Moreover, as described previously for other ETT-protein interactions, the sensitivity was specific to IAA as other auxinic compounds tested did not show this effect (Figure 3-figure supplement 2). Henceforth, ‘auxin’ will refer to IAA unless stated otherwise. These data suggest that in conditions with low auxin levels, ETT can interact with TPL/TPR proteins to repress the expression of target genes. An increase in cellular auxin causes ETT to bind auxin thereby undergoing a conformational change that abolishes interaction with TPL/TPR co-repressors.

TPL was originally identified as a key factor involved in setting up the apical-basal growth axis during embryo development^21, 22^. Large-scale interaction studies suggest that the five Arabidopsis TPL/TPRs have roles throughout plant development^20, 23^. Whilst ETT has been implicated in a wide array of developmental processes^24–27^, the most dramatic phenotypes of *ett* loss-of-function mutants are observed during gynoecium development^13, 28, 29^. In accordance with this, *ETT* is highly expressed in the gynoecium (Fig. 4a)^4^. We produced reporter lines of *TPL*, *TPR2* and *TPR4* promoters fused to the *GUS* gene to test if they overlap with *ETT* expression in the gynoecium. Both *pTPL:GUS* and *pTPR2:GUS* exhibited strong expression in the apical part of the gynoecium where *ETT* is also expressed, while no *pTPR4:GUS* expression was observed (Fig. 4a-d). Single loss-of-function mutants in *TPL* and *TPR2* do not show any abnormal phenotypes during gynoecium development. However, the *tpl tpr2* double mutant has defects in the development of the apical gynoecium similar to *ett* mutants (Fig. 4e-g) demonstrating that *TPL* and *TPR2* function redundantly in gynoecium development. Together with the protein interaction data and the overlapping expression patterns, these results suggest that ETT and TPL/TPR2 cooperate to regulate gynoecium development.

**Figure 4.**
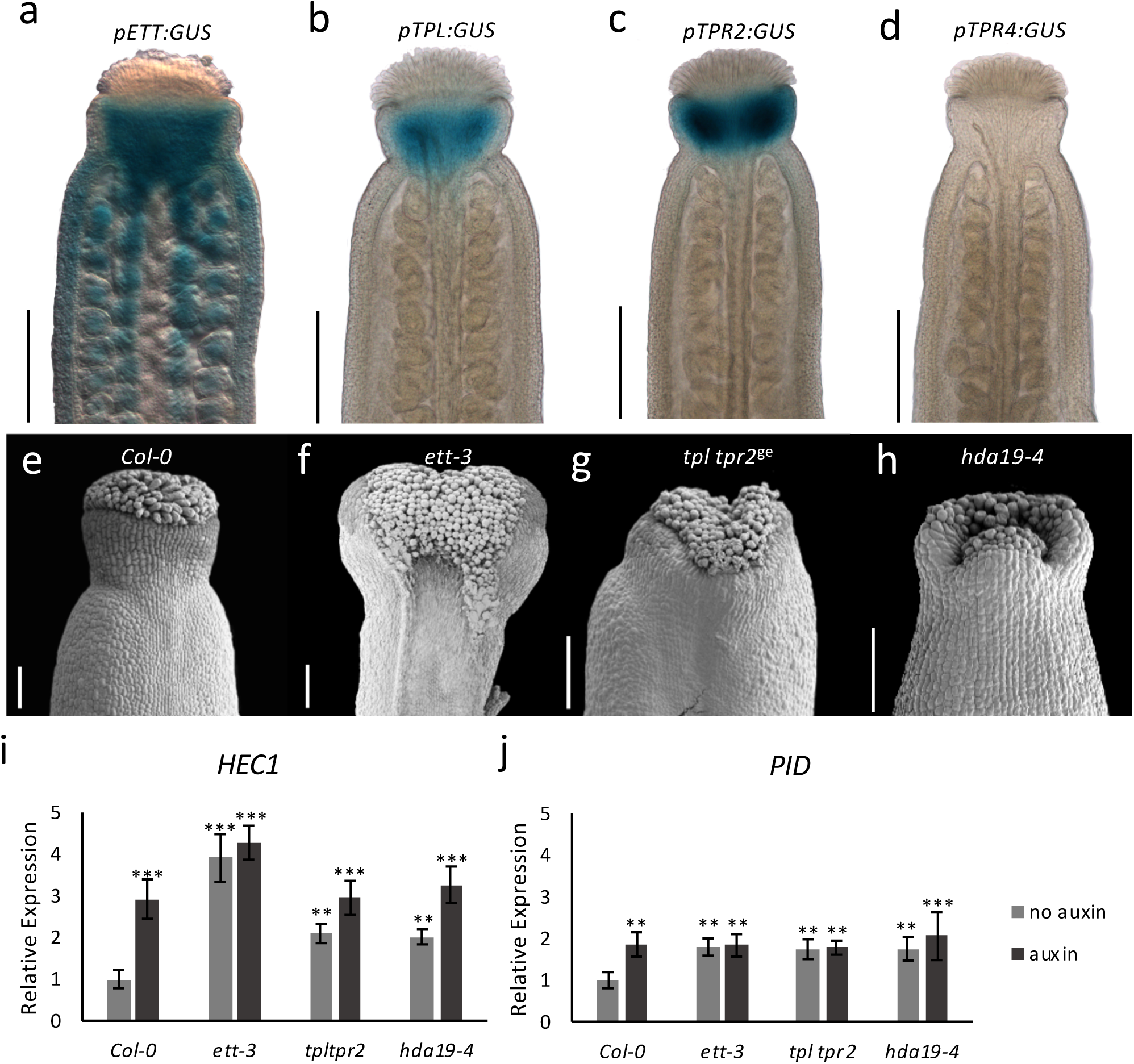
ETT, TPL/TPR2 and HDA19 co-operatively regulate gene expression to facilitate gynoecium development. **a-d,** Promoter GUS expression analysis of *pETT:GUS* (**a**), *pTPL:GUS* (**b**), *pTPR2:GUS* (**c**) and *pTPR4:GUS* (**d**) revealed that *ETT*, *TPL* and *TPR2* but not *TPR4* are co-expressed in the Arabidopsis style. Scale bar = 300 μm. **e-h,** Gynoecium phenotypes of wild-type (**e**), *ett-3* (**f**) *tpl tpr2^ge^* (**g**) and *hda19-4* (**h**). Scale bar = 100 μm. **i, j,** *HEC1* (**i**) and *PID* (**j**) are constitutively mis-regulated in *ett-3*, *tpl tpr2^ge^* and *hda19-4* gynoecia. This misregulation is unaffected by treatment with 100 µM IAA. ***p-Values<0.0001; Shown are mean ± standard deviation of three biological replicates. Differences between untreated and IAA-treated mutants are not significant. See Figure 4-source data 1 for statistical analyses.

TPL was shown previously to recruit histone deacetylase, HDA19, during early Arabidopsis flower development to keep chromatin in a repressed state^20^. Moreover, HDA19 was also recently shown to participate in repression of the meristem identity gene, *SHOOT MERISTEMLESS* (*STM*)^30^. Here, our analysis of gynoecia from the *hda19-4* mutant demonstrate that HDA19 is also required for gynoecium development as the *hda19-4* mutant has strong style defects (Fig. 4h). In agreement with this, the *HDA19* gene was highly expressed in gynoecium tissue, whereas another member of the *HDA* gene family, *HDA6*, was not (Figure 4-figure supplement 1). Moreover, HDA19 recruitment likely involves *ETT*, since expression of the ETT target genes, *PID* and *HEC1*, are increased in the *tpl tpr2* and *hda19-4* mutants compared to wild type. Similar to the *ett* mutant, auxin treatments failed to further induce expression in these mutants (Fig. 4i,j). These observations suggest that ETT, TPL/TPR2 and HDA19 function in conjunction to control gene expression during gynoecium development.

To test the direct interaction of ETT, TPL and HDA19 on chromatin, we performed Chromatin-Immunoprecipitation (ChIP) using reporter lines expressing GFP fusion protein. Although only ETT is expected to bind DNA, ChIP followed by qPCR revealed that all three proteins associate with DNA elements in the same regions of the promoters of *PID* and *HEC1* (Fig. 5a). This supports a model in which ETT recruits TPL/TPR2 and HDA19 to ETT target loci to keep chromatin in a condensed state through histone deacetylation. When auxin levels increase, the ETT-TPL/TPR2 interaction is broken, presumably preventing HDA19 from deacetylating histones. To test this, we assayed for H3K27 acetylation, which is a substrate for HDA19. H3K27 acetylation increased in the absence of ETT and upon treatment with auxin. This occurred in the same regions of the *PID* and *HEC1* promoters where the proteins were found to associate (Fig. 5b,c). In agreement with ETT mediating the association of TPL/TPR and HDA19 with these regions, there was no further increase of acetylation in the *ett-3* mutant upon treatment with auxin (Fig. 5b,c).

**Figure 5.**
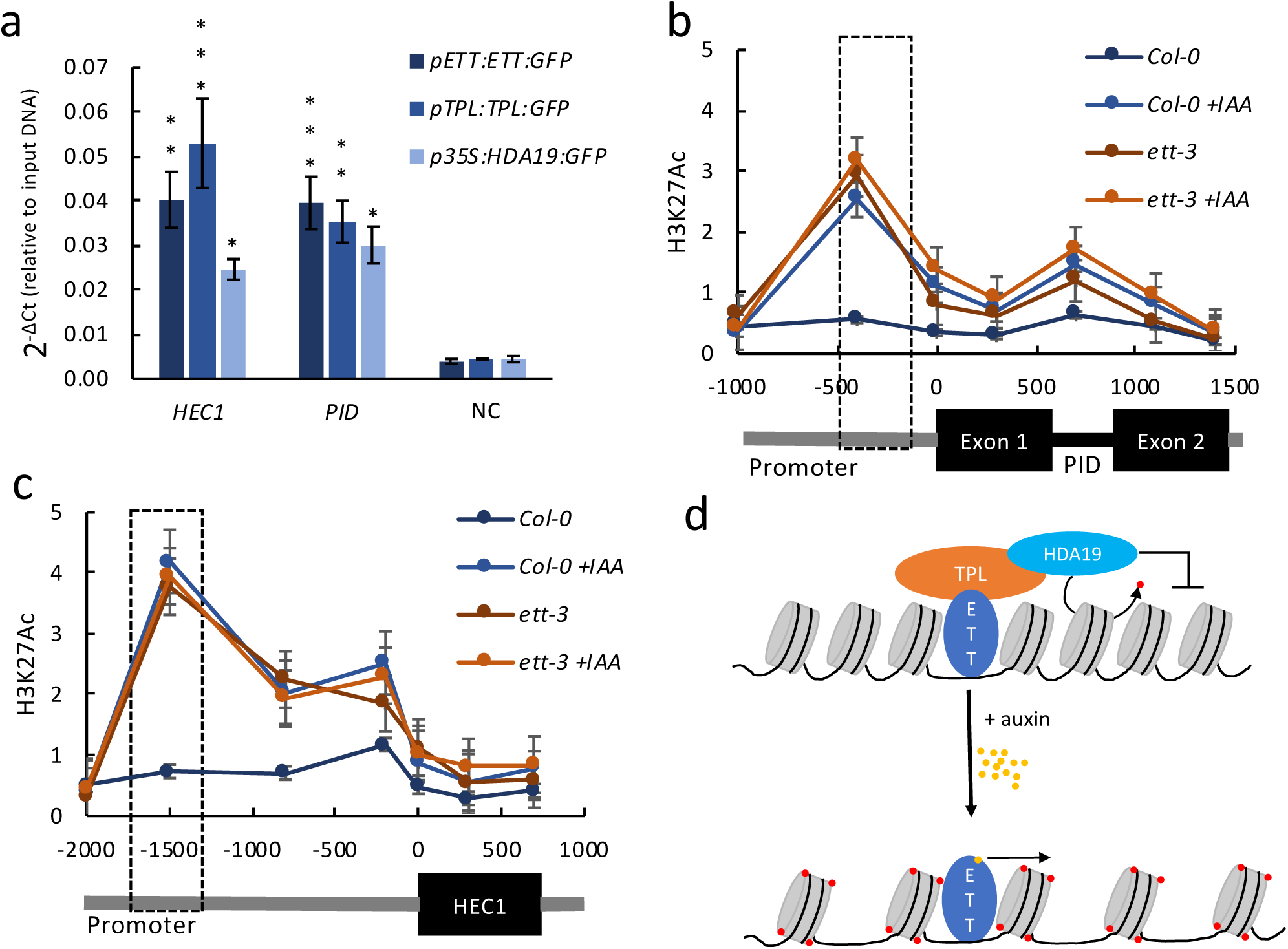
ETT, TPL and HDA19 co-operatively regulate *HEC1* and *PID* by modulation of chromatin acetylation. **A,** Chromatin immunoprecipitation (ChIP) shows ETT, TPL and HDA19 binding to conserved regions of *HEC1* and *PID* loci. *WUS* served as negative control. **b, c,** H3K27ac accumulation (from ChIP analysis) along the *HEC1* (**b**) and *PID* (**c**) loci in wild-type (*Col-0*) and *ett-3* plants ± treatment with 100 µM IAA. Numbers on the x axes are distances to the Transcription Start Site (TSS). The schematic of the loci is shown below each panel. Dashed boxes represent ETT binding regions. **d,** Schematic model illustrating alternative TIR1/AFB independent auxin signaling. Under low auxin conditions an ETT-TPL-HDA19 complex binds to ETT-target genes keeping their chromatin environments repressed, through de-acetylation. High nuclear auxin concentrations abolish the ETT-TPL-HDA19 complex through direct ETT-auxin interaction. This leads to an accumulation of histone acetylation and up-regulation of ETT-target genes. Values in **a**, **b** and **c** are means ± standard deviation of three biological replicates. See Figure 5-source data 1 for statistical analyses.

The data presented in this paper provide molecular insight as to how auxin levels are translated into changes in gene expression of ETT target genes. Our data lead to a model in which low levels of auxin maintain ETT associations with TPL/TPR2 to repress gene expression via H3K27 deacetylation. As auxin levels increase, TPL/TPR2 (and hence HDA19) disassociate from ETT, promoting H3K27 acetylation (Fig. 5d). This model molecularly underpins the published association between auxin dynamics and *PID* expression at the gynoecium apex where *PID* is repressed at early stages of development to allow symmetry transition, but subsequently de-repressed as auxin levels rise to facilitate polar auxin transport^4, 31^.

The direct binding of auxin allows ETT to switch the chromatin locally between repressive and de-repressive states, whilst other ARFs have been categorised as either repressors or activators^8^. The effect of auxin is therefore instantly reversible, making it possible to switch between states immediately in response to changes in auxin levels. This feature, which is reminiscent of animal hormonal signalling pathways such as the Thyroid Hormone and Wnt/ß-catenin pathways^1, 2^, may be particularly important in controlling changes in tissue polarity during plant organogenesis as observed in the Arabidopsis gynoecium^31^.

The identification of a direct auxin-ETT interaction to control gene expression adds an additional layer of complexity to auxin biology, which contributes towards explaining how auxin imparts its effect on highly diverse processes throughout plant development. In a wider context, this work also opens for the exciting possibility that direct transcription factor-ligand interactions is a general feature in the control of gene expression in plants as found in animals.

## MATERIAL AND METHODS

### Plant materials and treatments

Plants were grown in soil at 22 °C in long day conditions (16hrs day/8 hrs dark). All mutations were in the *Col-0* background. Mutant alleles described before include e*tt-3*^4, 13^*, hda19-4* (SALK_139443)^32^, *pETT:GUS*^33^, *pETT:ETT-GFP* in *ett-3*^4^, *pTPL:TPL:GFP*^34^, *p35S:HDA19:GFP*^34^, *pTIR1:ccvTIR1* in *tir1-1 afb2-3*^19^ and *tir1-1 afb2-3 afb3-4*^35^.The *tpl* mutant (SALK_034518C) was obtained from the European Arabidopsis Stock Centre. For both expression and ChIP analysis, auxin treatments were applied by spraying bolting *Col-0* and *ett-3* inflorescences with a solution containing 100 µM IAA (Sigma) or cvxIAA and 0.015% Silwet L-77 (De Sangosse Ltd.). Treated samples were returned to the growth room and incubated for two hours.

### Expression analysis

Quantitative Real time PCR (qRT-PCR) was used for expression analysis. RNA was extracted from floral buds using the RNeasy mini kit (Qiagen). Using the SuperScript™ IV First-Strand Synthesis kit (ThermoFisher), cDNA was synthesised from 1 µg of total RNA. Subsequently, qRT-PCR was carried out using SYBR Green JumpStart Taq ReadyMix (Sigma) using the appropriate primers (Figure 2-source data 1). Relative expression values were determined using the 2^-ΔΔCt^ method^36^. Data were normalised to *POLYUBIQUITIN 10* (*UBQ10/*AT4G05320) expression.

### ETT protein analysis by alignment

Published *ETT* sequences of 22 Angiosperm species were retrieved from Phytozome version 12^37^. Nucleotide sequences were translated and aligned using MUSCLE in Geneious version 6.1.8^38^. The EAR domain was extracted as a sequence logo (Fig. 3a; Figure 3-figure supplement 1).

### Generation of the *tpl tpr2* CRISPR mutant

The *tpl tpr2^ge^* mutant was generated using CRISPR/Cas9 technology by a method previously described^39^. Briefly, for the construction of the RNA-guided genome-editing plasmid, DNA sequences encoding the gRNA adjacent to the PAM sequences were designed to target two specific sites in *TPR2* (AT3G16830). DNA-oligonucleotides (Figure 2-source data 1) containing the specific gRNA sequence were synthesised and used to amplify the full gRNA from a template plasmid (AddGene #46966). Using Golden Gate cloning^40^ each gRNA was then recombined in a L1 vector downstream of U6 promoter^39^. Finally, the resulting gRNA plasmids were then recombined with a L1 construct containing *pYAO:Cas9_3:E9t*^39^ (kindly provided by Jonathan Jones) and a L1 construct containing Fast-Red selection marker (AddGene #117499) into a L2 binary vector (AddGene #112207).

The construct was transformed into *Agrobacterium tumefaciens* strain GV3101 by electroporation, followed by plant transformation by floral dip into the *tpl* single mutant^41^. Transgenic T0 seeds appear red under UV light and were selected under a Leica M205FA stereo microscope. T0 plants were genotyped using PCR and the *TPR2* locus sequenced (Oligonucleotides in Figure 2-source data 1). Genome edited plants were selected and the next generation grown (T1). Seeds of this generation were segregating in a 3:1 ratio for the transgene. Transgene negative plants were selected and grown on soil. To find homozygous mutations T1 plants were again genotyped. The T2 generation was again checked for the absence of the transgene.

### Protein interaction

For Yeast-two-Hybrid (Y2H) assays coding sequences were cloned into pDONR207 and recombined into the pGDAT7 and pGBKT7 (Clontech). Using the co-transformation techniques^41^ these constructs were transformed into the AH109 strain (Clontech).

Transformations were selected on Yeast Selection Medium (YSD) lacking Tryptophan (W) and Leucine (L) at 28°C for 3-4 days. Transformed yeast cells were serially diluted (10^0^, 10^1^, 10^2^ and 10^4^) and dotted on YSD medium lacking Tryptophan (W), Leucin (L), adenine (A) and Histidine (H) to test for interaction. To examine interaction strength 3-amino-1,2,4-triazole (3-AT) was supplemented to the YSD (-W-L-A-H) medium with different concentrations (0, 5, 10 mM). To determine the effect of auxinic compounds on the protein-protein interactions benzoic acid (BA), IAA, NAA and 2,4D (all Sigma) were dissolved in ethanol and added directly to the medium at the desired concentrations. Pictures were taken after 3 days of growth at 28°C.

For the β-Galactosidase assay transgenic yeast was grown in liquid YSD (-W-L) medium supplemented with/-out 100 µM IAA or NAA, to an OD_600_ of 0.5. The cells were then harvested and lysed using 150 µL Buffer Z with β-mercaptoethanol (100 mM Phosphate buffer pH 7, 10 mM KCl, 1mM Mg_2_SO_4_, β-mercaptoethanol 50 mM), 50 µL chloroform and 20 µL of 0.1% SDS. After lysis, the sample was incubated with 700 µL pre-warmed ONPG solution (1mg/mL ONPG (o-Nitrophenyl-β-D-Galactopyranoside, Sigma) prepared in Buffer Z without β-mercaptoethanol at 28°C until a yellow colour developed in the samples without auxin treatment. After stopping all reactions (using 500 µL Na_2_CO_3_) the supernatant was collected and OD_405_ determined. The β-Galactosidase activity was calculated as follows: (A405*1000)/(A600*min*mL)

For co-immunoprecipitation, ETT-FLAG was generated using Golden Gate cloning^39^ by recombining a previously described L0 clone for ETT with a 35S promoter (AddGene #50266), a C-terminal 3xFLAG epitope (AddGene #50308) and a Nos-terminator (AddGene #50266) into a L1 vector (AddGene #48000). The pGWB14 TPL-HA construct was provided by Salomé Prat and has been used in previous studies^42^. The epitope-tagged proteins were transiently expressed in four-week-old *N. benthamiana* leaves for two days. Co-immunoprecipitation was performed as described previously^43^. After harvest, 1 g of fresh leaf tissue was ground in liquid nitrogen. The powder was homogenised for 30 min in two volumes of extraction buffer (10% glycerol, 25 mM Tris-HCl pH 7.5, 1 mM EDTA, 150 mM NaCl, 0.15% NP-40, 1mM PMSF, 10 mM DTT, 2% Polyvinylporrolidone, 1x cOmplete Mini tablets EDTA-free Protease Inhibitor Cocktail (Roche). The homogenised samples were cleared by centrifugation at 14,000 xg for 10 min and cleared lysates were incubated for 2h with 20 µl anti-FLAG M2 magnetic beads (SIGMA-ALDRICH, M8823; lot: SLB2419). The beads were washed five times with IP buffer (10% glycerol, 25 mM Tris-HCl pH 7.5, 1 mM EDTA, 150 mM NaCl, 0.15% NP-40, 1 mM PMSF, 1 mM DTT, 1x cOmplete Mini tablets EDTA-free Protease Inhibitor Cocktail (Roche)) and proteins were eluted by adding 80 µl 2x SDS loading buffer followed by an incubation at 95 °C for 10 min. To examine auxin sensitivity 4 g of fresh leaf tissue was collected, ground in liquid nitrogen and protein was extracted. The lysate was then divided according to the number of treatments. The desired concentration of IAA or NAA was added to each of the cleared lysates before the anti-FLAG M2 magnetic beads were added. IAA or NAA at the desired concentration was also supplemented to the IP buffer during the washes. The eluates were analysed by western blot using an anti-FLAG antibody (M2, Abcam, ab49763, Lot: GR3207401-3) or an anti-HA antibody (Abcam, ab173826, Lot: GR3255539-1). Both antibodies were used as 1:10000 dilutions. The antibodies were validated by the manufacturer.

### Scanning electron microscopy

Whole inflorescences of *Col-0*, *ett-3*, *tpl tpr2^ge^* and *hda19-4* were fixed overnight in FAA (3.7% formaldehyde, 5% glacial acetic acid, 50% ethanol) and dehydrated through an ethanol series (70% to 100%) as described previously^31^. The samples were then critical point-dried, gynoecia dissected and mounted. After gold coating samples were examined with a Zeiss Supra 55VP Field Emission Scanning electron microscope using an acceleration voltage of 3 kV.

### *TPL*, *TPR2* and *TPR4* reporter lines

For the construction of the promoter:GUS reporter plasmids of *TPL*, *TPR2* and *TPR4*, 2.5 kb of promoter sequences were isolated from genomic DNA and inserted upstream of the ß-glucoronidase gene of pCambia1301 vectors using the In-Fusion Cloning Recombinase kit (Clontech). The constructs were transformed into *Agrobacterium tumefaciens* strain GV3101 by electroporation, followed by plant transformation by floral dip into *Col-0*^41^.

The GUS histochemical assay was performed in at least three individual lines per construct. Inflorescences of each GUS line were pre-treated with ice cold acetone for 1h at −20°C and washed two times for 5 minutes with 100 mM sodium phosphate buffer followed by one wash with sodium phosphate buffer containing 1 mM K_3_Fe(CN)_6_ and 1 mM K_4_Fe(CN)_6_ (both Sigma) at room temperature. Subsequently, samples were vacuum infiltrated for 5 minutes with X-Gluc solution (100 mM sodium phosphate buffer, 10 mM EDTA, 0.5 mM K_3_Fe(CN)_6_, 3 mM K_4_Fe(CN)_6_, 0.1% Triton X100) containing 1 mg/ml of ß-glucoronidase substrate X-gluc (5-bromo-4-chloro-3-indolylglucuronide, Melford) and incubated at 37°C. *pTPL:GUS* were incubated for 20 minutes and *pTPR2:GUS* lines for 45 minutes to prevent overstaining. *pTPR4:GUS* lines were incubated for 16h. After staining, the samples were washed in 70% ethanol until chlorophyll was completely removed. Gynoecia were dissected and mounted in chloral hydrate (Sigma). Samples were analysed using a Leica DM6000 light microscope.

### Chromatin Immunoprecipitation

Transcription factor ChIP was performed in triplicate using the *pETT:ETT:GFP*, *pTPL:TPL:GFP* and *p35S:HDA19-GFP* lines and data analysed as described previously^44^. Additionally, a *WUS* promoter fragment was used as a negative control for ETT binding^45^. IP was conducted using the anti-GFP antibody (Roche, 11814460001, Lot: 19958500) and Pierce Protein G magnetic beads (ThermoFisher, 88847, Lot: SI253639) were used for IP. Histone acetylation ChIP was carried out and data were analysed as described previously^46^. The experiment was carried out in triplicate using 3 g auxin-treated or untreated *Col-0* or *ett-3* inflorescent tissue. The antibodies used for IP were anti-H3K27ac antibodies (Abcam, ab4729, Lot: GR3231937-1) and anti-H3 (Abcam, ab1791, Lot: GR310541-1). All antibodies were validated by the manufacturers.

In all ChIP experiments, DNA enrichment was quantified using quantitative PCR (qPCR) with the appropriate primers (Supplementary Data). In case of H3K27ac, *ACTIN* was used as an internal control and the data represented as ratio of (H3K27ac at *HEC1* or *PID* divided by H3 at *HEC1or PID*) to H3K27ac at *ACT* divided by H3 at *ACT*).

### Statistical analyses and replication

In all graphs error bars represent the standard deviation of the mean for all numerical values. QRT-PCR and ChIP experiments have been carried out at least in triplicate. The data presented here show an average of three replicates. For qRT-PCR data were analysed using one-way ANOVA with post-hoc Tukey multiple comparison test. ChIP_qPCR_ data were analysed using two-way ANOVA with post hoc Bonferroni multiple comparison test. All output of statistical tests can be found in the source data files. All statistical tests were carried out using GraphPad Prism Version 5.04 (La Jolla California USA, www.graphpad.com).

### Protein production

The ES domain, ES^388–594^, protein was isotopically labelled in preparation for NMR analysis. The ES domain was expressed for as a fusion protein with a 6x Histidine tag in minimal media with ^15^N ammonium chloride. The ^15^N isotope labelling of the expressed protein involved a 125-fold dilution of cell culture in enriched growth media into minimal media with ^15^N ammonium chloride and grown for 16 hours (37 °C / 200 rpm); followed by a further 40-fold dilution into minimal media for the final period of cell growth and protein expression (induced with L-arabinose 0.2 % w/v / 18 °C / 200 rpm and grown for a further 12 hours). The fusion protein was isolated from soluble cell lysate by Co-NTA affinity chromatography with two His-Trap 1 mL TALON Crude columns (GE Healthcare Life Sciences, 28953766). Chromatography buffers contained sodium phosphate 20 mM pH 8.0, NaCl 500 mM and either no-imidazole or 500 mM imidazole for wash and elution buffers respectively. The majority of the non-specifically bound protein was removed by passing 20 mL of the wash buffer through the columns. The protein eluted on a gradient of increasing imidazole concentration of up to 30% elution buffer over 20 mL.

### HSQC NMR

The ES domain, ES^388-594^, protein was analysed by NMR at 5°C under reducing conditions (DTT 10 mM), buffered at pH 8.0 (Tris 20 mM). ^1^H-^15^N HSQC was performed at 950 MHz, TCI probe, Bruker following the parameters described in Figure 1-figure supplement 1.

### Isothermal titration calorimetry (ITC)

ITC was carried out on a MicroCal PEAQ-ITC (Malvern) at 25°C in a Buffer A (sodium phosphate 20 mM, pH 8.0; NaCl 500 mM). Ligand (2 mM IAA) was injected (19 × 4.0 μl) at 150-s intervals into the stirred (500 rpm) calorimeter cell (volume 270 μl) containing 50 µM ES^388-594^ protein. Titration of Buffer A into 50 µM ES^388-594^ protein and IAA (2 mM) into Buffer A served as negative controls. Measurements of the binding affinity of all the titration data were analysed using the MicroCal Software (Malvern).

### Accessions

ETT, AT2G33860; TPL, AT1G15750; TPR1, AT1G80490, TPR2, AT3G16830; TPR3, AT5G27030; TPR4, AT3G15880; HDA6, AT5G63110; HDA19, AT4G38130; HEC1, AT5G67060; PID, AT2G34650; WUS, AT2G17950.

## Acknowledgements

We are grateful to Yuli Ding, Yang Dong, Emilie Knight, Bhavani Natarajan, Mikhaela Neequaye, Nicola Stacey, Sophia Stavnstrup, Billy Tasker-Brown for critical comments on the manuscript, to Keiko Torii and Shinya Hagihara for the ccvTIR1 line and cvxIAA ligand, to Rebecca Mosher for assistance with the phylogenetic analysis of ETT protein sequences, to Thomas Laux for *TPL::TPL-GFP* and *35S::HDA19-GFP* lines and to Salomé Prat for *35S::TPL-HA* construct. We acknowledge Norwich Research Park Bioimaging for skillful assistance and the NMR facility in the Astbury Centre, Faculty of Biological Sciences for access to the 950 MHz and 600 MHz spectrometers funded by the University of Leeds. We thank Arnout Kalverda for assistance with analysing the NMR data.

## Author Contributions

A.K. and L.Ø. conceived the experiments. A.K., S.R.H. and H.M.M. performed the experiments. A.K., S.R.H., H.M.M., S.K. and L.Ø. analysed the data. A.K. and L.Ø. wrote the manuscript and S.R.H., H.M.M. and S.K. commented on it. All authors read and approved the manuscript.

## Ethics

Human subjects: No; Animal subjects: No

## Dual-use research

No

## Permissions

Have you reproduced or modified any part of an article that has been previously published or submitted to another journal?

No

---

**Figure 1-figure supplement 1.**
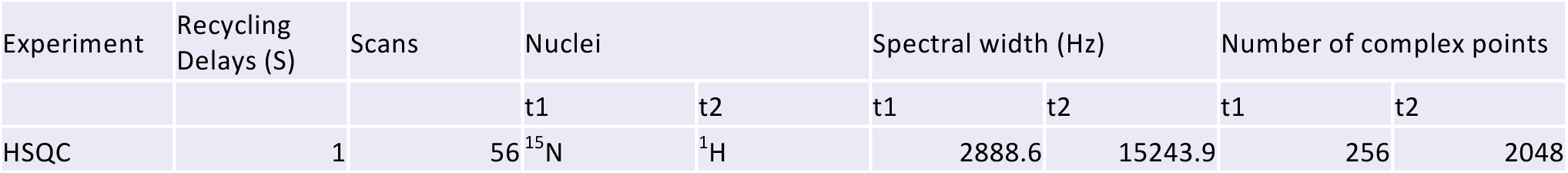
Parameters for HSQC NMR experiment.

**Figure 2-figure supplement 1.**
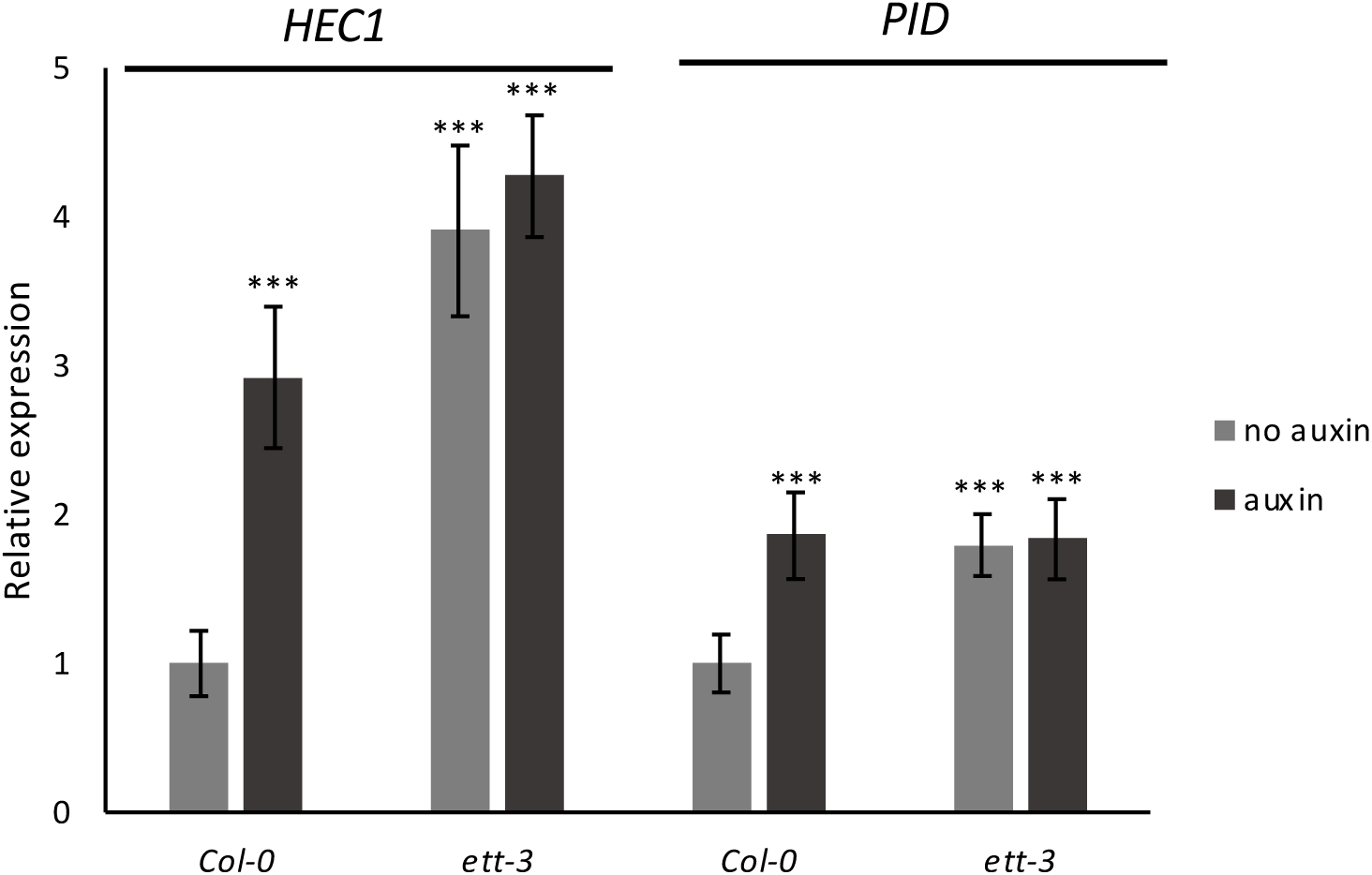
Expression of *HEC1* and *PID* in Col-0 and *ett-3*. In wild-type gynoecia *HEC1* and *PID* are up-regulated upon auxin treatment while both genes are constitutively up-regulated in *ett-3*. Treatment with 100 µM IAA does not affect *HEC1* and *PID* expression in the *ett-3* mutant suggesting that ETT acts as a transcriptional repressor. Asterisks indicate significant change upon auxin treatment compared to untreated *Col-0* (*** indicating p < 0.0001). Shown are mean ± standard deviation of three biological replicates. See Figure 2-source data 1 for statistical analyses.

**Figure 3-figure supplement 1.**
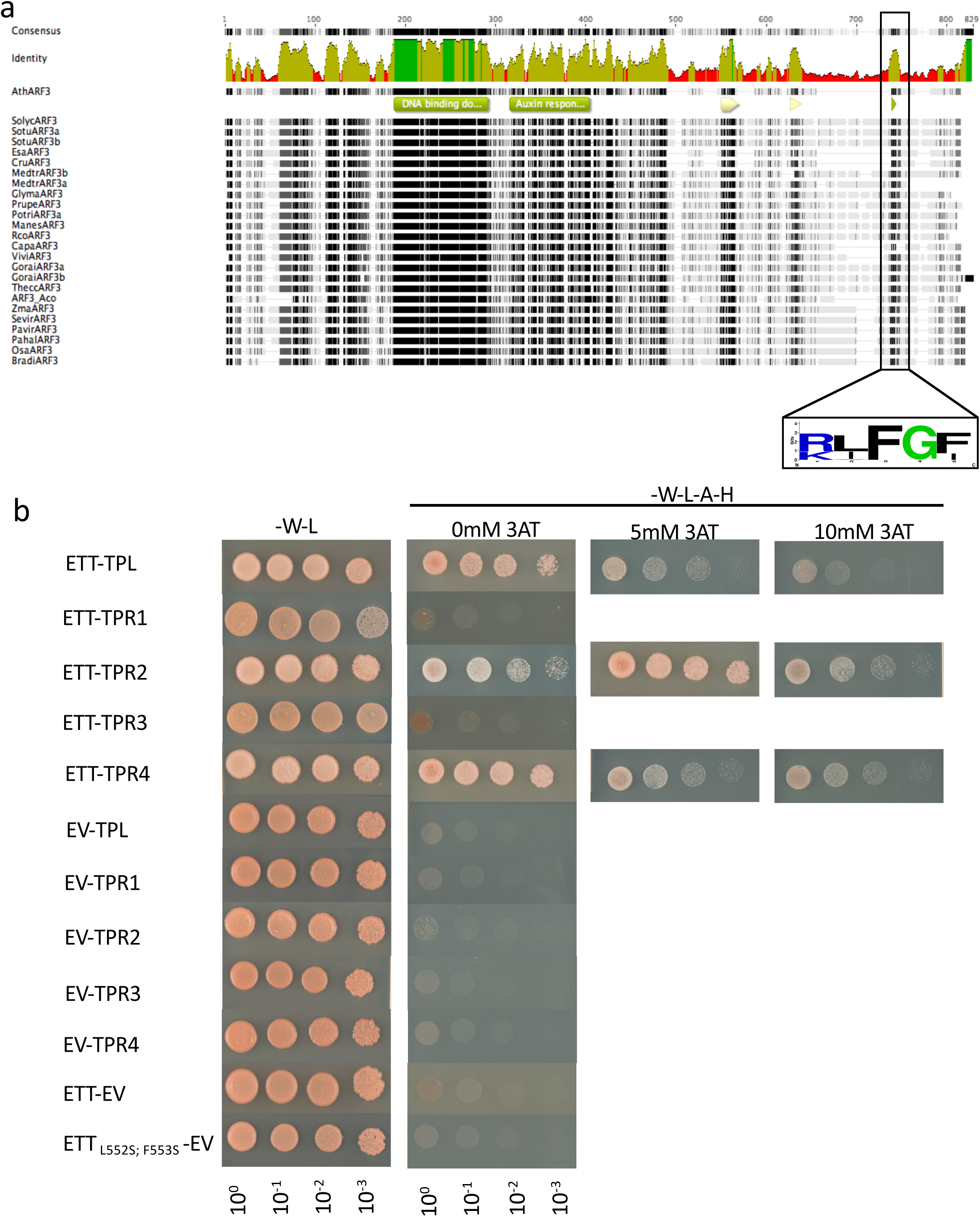
ETT can interact with several members of the TPL/TPR co-repressor family through a conserved EAR-like motif. **a,** Alignment of ETT protein sequences of 22 species identified a conserved repressive motif (RLFGF) at its c-terminal domain. **b,** ETT interacts with several members of the TPL/TPR co-repressor family in Y2H. Additionally, controls for Fig. 2 are shown.

**Figure 3-figure supplement 2.**
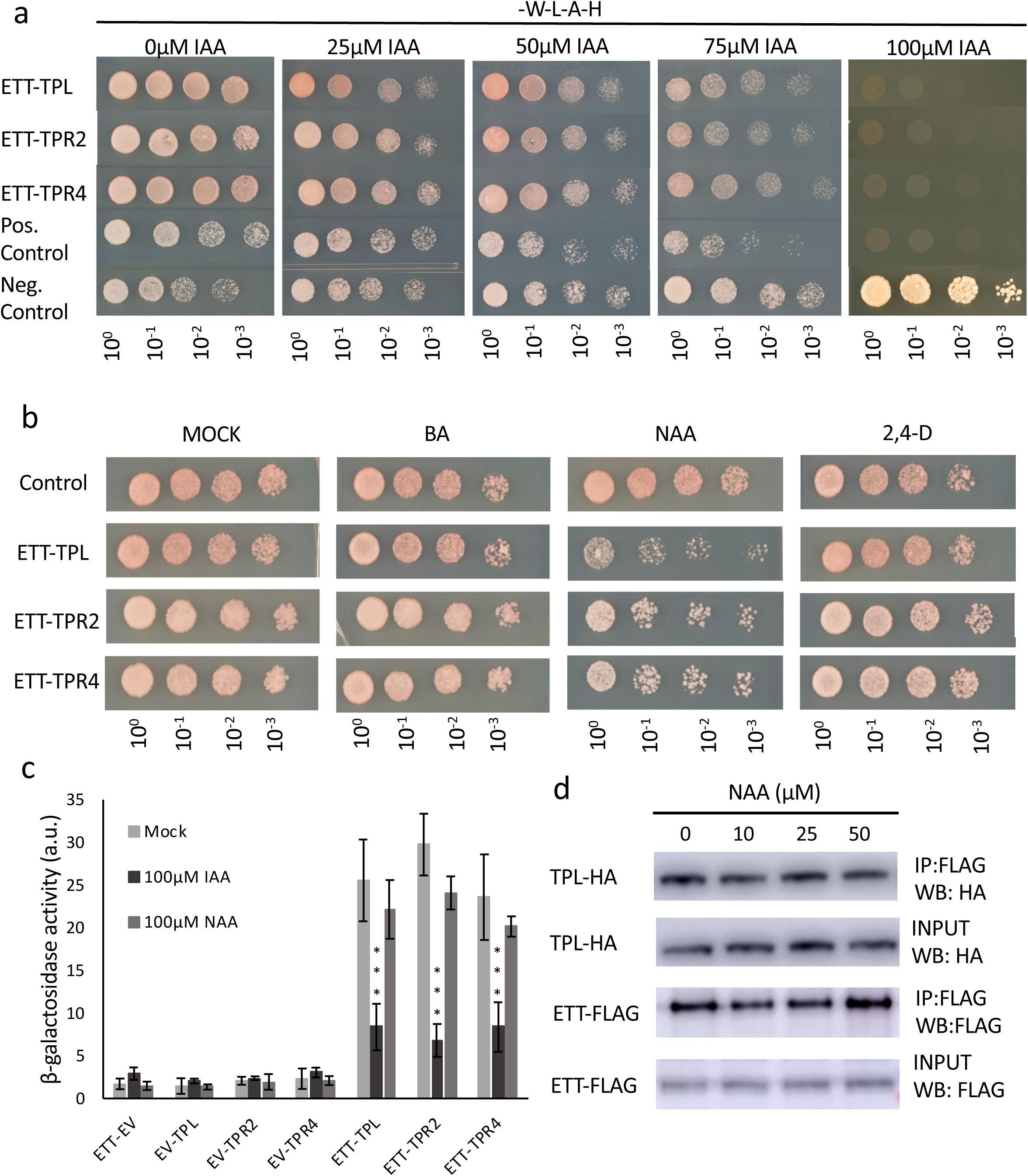
Interaction between ETT and TPL, TPR2 and TPR4 is auxin-sensitive and specific to IAA. **a,** In Y2H increasing concentrations of IAA lead to reduction of yeast growth abolishing the interaction between ETT and its partners. The interactions are, therefore, auxin-sensitive. **b,** Y2H to test specificity of auxin-sensitivity using benzoic acid (BA), NAA, and 2,4D in a yeast growth assay. The data suggest that the auxin-sensitivity observed in (**a**) is IAA-specific. **c,** Y2H based ONPG assay measuring the β-galactosidase activity as a measure of interaction strength. **d,** Co-IP experiments show that the interaction between ETT and TPL cannot be disrupted by NAA. The data support that the ETT TPL/TPR interactions are sensitive to IAA but not to NAA. ***p <0.0001; Shown are mean ± standard deviation of three biological replicates. See Figure 3-source data 1 for statistical analyses.

**Figure 3-figure supplement 3.**
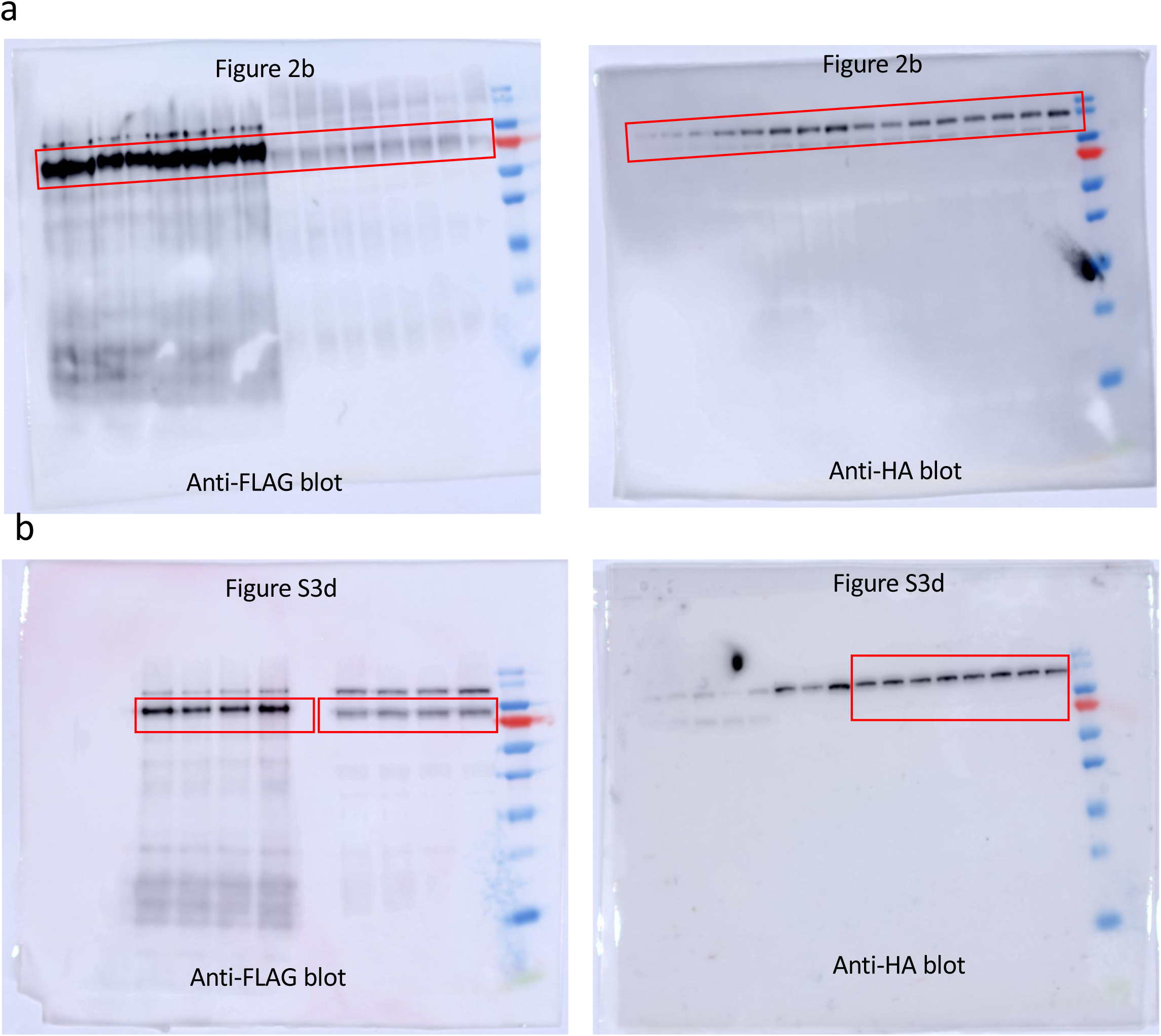
Original western blot images. The red boxes indicate the areas used in Figure 3b and Figure 3-figure supplement 2d.

**Figure 4-figure supplement 1.**
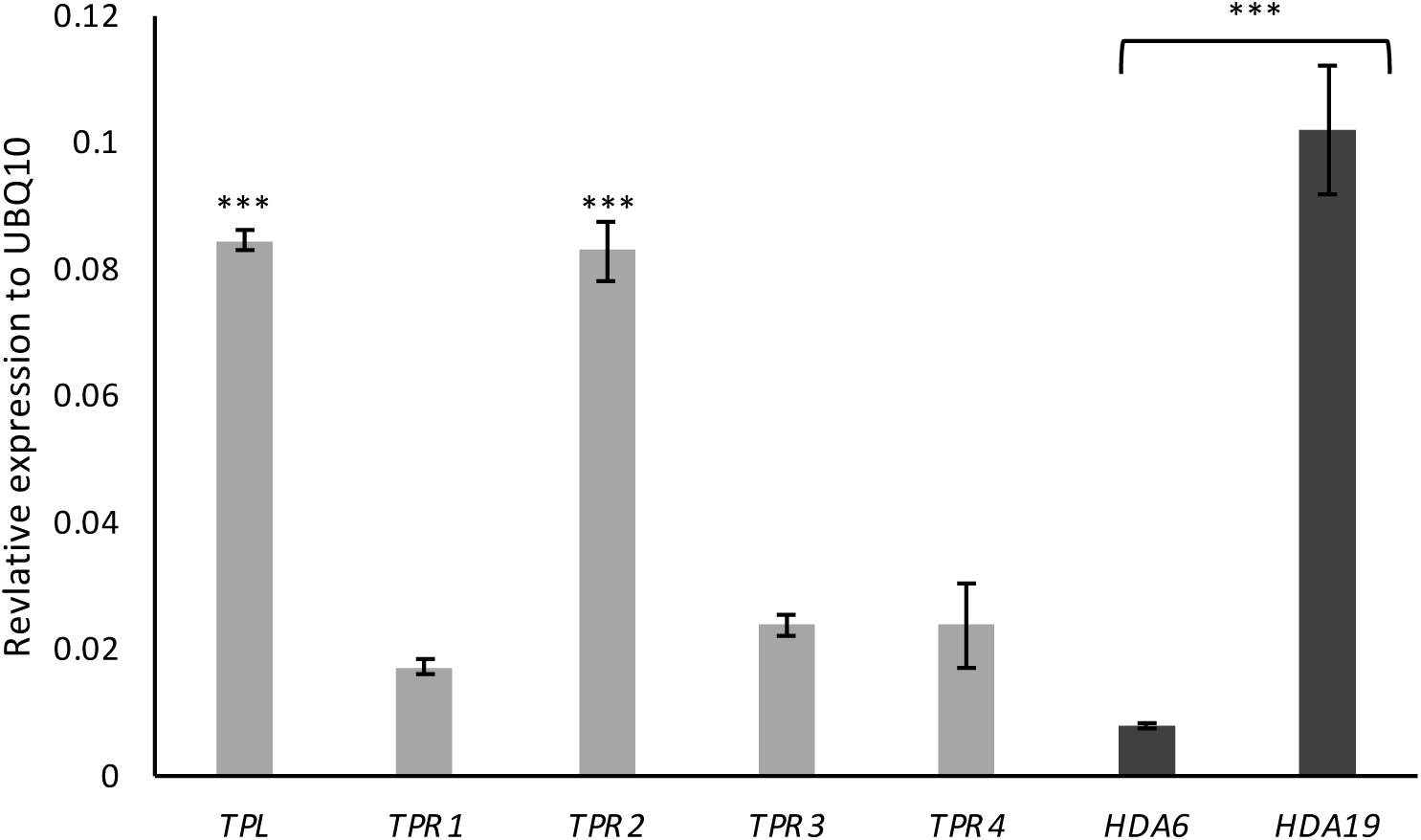
Expression of *TPL*, *TPRs* and *HDAs* genes in the gynoecium. Expression analysis using qRT-PCR in wild-type gynoecia showed that *TPL* and *TPR2* are more strongly expressed than *TPR1*,*3* and *4*. Likewise, *HDA19* exhibits higher expression compared to *HDA6*. ***p-Values<0.0001; Shown are mean ± standard deviation of three biological replicates. See Figure 4-source data 1 for statistical analyses.

## Source data

**Figure 1-source data 1. Parameters for HSQC NMR.**

**Figure 2-source data 1. Output of statistical tests**

**Figure 3-source data 1. Output of statistical tests**

**Figure 4-source data 1. Output of statistical tests**

**Figure 5-source data 1. Output of statistical tests**

**Supplementary Data. Oligonucleotides used in this study.**

